# The PNUTS-PP1 axis regulates endothelial aging and barrier function via SEMA3B suppression

**DOI:** 10.1101/2020.08.10.243170

**Authors:** Noelia Lozano-Vidal, Laura Stanicek, Diewertje I. Bink, Veerle Kremer, Alyson W. MacInnes, Stefanie Dimmeler, Reinier A. Boon

## Abstract

Age-related diseases pose great challenges to health care systems worldwide. During aging, endothelial senescence increases the risk for cardiovascular disease. Recently, it was described that Phosphatase 1 Nuclear Targeting Subunit (PNUTS) has a central role in cardiomyocyte aging and homeostasis. Here, we determined the role of PNUTS in endothelial cell aging. We confirmed that PNUTS is repressed in senescent endothelial cells (ECs). Moreover, PNUTS silencing elicits several of the hallmarks of endothelial aging: senescence, reduced angiogenesis and loss of barrier function. To validate our findings in vivo, we generated an endothelial-specific inducible PNUTS-deficient mouse line (Cdh5-CreERT2;PNUTS^fl/fl^), termed PNUTS^EC-KO^. Two weeks after PNUTS deletion, PNUTS^EC-KO^ mice presented severe multiorgan failure and vascular leakage. We showed that the PNUTS binding motif for protein phosphatase 1 (PP1) is essential to maintain endothelial barrier function. Transcriptomic analysis of PNUTS-silenced HUVECs and lungs of PNUTS^EC-KO^ mice revealed that the PNUTS-PP1 axis tightly regulates the expression of semaphorin 3B (SEMA3B). Indeed, silencing of SEMA3B completely restored barrier function after PNUTS loss-of-function. These results reveal a pivotal role for PNUTS in endothelial homeostasis through a PP1-SEMA3B downstream pathway that provides a potential target against the effects of aging in ECs.

## INTRODUCTION

Cellular aging is characterized by hallmarks such as epigenetic changes, genomic instability, mitochondrial stress and cellular senescence that promote a certain number of processes leading to physiological dysfunction and multiple pathologies [1]. The progressive aging of the population in Western countries has led to an increase of age-related diseases, a new health care challenge [2]. Of those, cardiovascular diseases (CVD) have the highest prevalence in the elderly, both in women and men. Therefore, aging is the principal risk factor for cardiovascular diseases [3].

The vascular endothelium serves as a barrier between the blood and the surrounding tissues, and has other critical functions for homeostasis, such as maintaining vascular tone and controling platelet activity, leukocyte adhesion and angiogenesis [4]. The molecular cues of aging induce endothelial dysfunction, a broad term that includes several characteristics, such as reduced vasodilatory and antithrombotic properties and an increase in the expression of pro-oxidant and pro-inflammatory genes [5]. Aged endothelial cells present lower proliferative and migration capacity, higher sensitivity to apoptotic signals and cellular senescence [6]. As a result, endothelial aging is correlated with impaired angiogenesis, reduced capillary density, increased vascular permeability and a pro-thrombotic phenotype in vascular beds, all leading to CVD [7].

Vascular permeability is controlled by different mechanisms involving intrinsic and extrinsic factors. Among these, there are changes in the composition of intercellular adherens and tight junctions and focal focal adhesions [8] [9], the contractile function of the actin-myosin cytoskeleton [10], paracrine signalling [11] and variations in calcium transients [12]. An important regulator of vascular integrity is the family of axon guidance proteins, specifically the semaphorin subfamily. Initially identified as determinants of axon growth and polarization [13], semaphorins are shown now to have an important role in regulating vascular patterning, endothelial junctions, angiogenesis and barrier function [14], [15], [16], [17].

Phosphatase 1 Nuclear Targeting Subunit (PNUTS, *PPP1R10*) is a nuclear protein described to act as a binding platform for the PP1 phosphatase complex [18]. PNUTS is implicated in cell cycle, cancer cell proliferation and protection against DNA damage response and apoptosis [19] [20] [21]. Recently, we showed that PNUTS expression is markedly repressed in the heart during aging and CVD, and the mechanism by which PNUTS downregulation induces post-ischemic heart failure includes telomere attrition in cardiomyocytes, a hallmark of aging [22]. PNUTS protein contains multiple binding domains, serving as partner for several targets, as PP1, TRF2, PTEN and nucleic acids, which determine PNUTS function in different contexts [23] [24] [25]. The PNUTS-PP1 complex is formed by interaction through the canonical RxVF motif. By binding to PP1, PNUTS is implicated in the dephosphorylation and activity of several PP1 targets, such as RNA polymerase II [26], Myc [27], Rb [28] and Spt5 [29]. However, whether PNUTS plays a role in endothelial aging is not known.

Here, we show that PNUTS is an aging-regulated gene in the endothelium, and that PNUTS depletion mimics an aging endothelial phenotype. Also, we describe for the first time an endothelial-specific PNUTS knock-out animal model which presents severe endothelial dysfunction similar to PNUTS depletion in cultured human endothelial cells. Particularly, endothelial PNUTS loss triggers a dramatic increase in endothelial permeability, but has no effect on adherens junction composition. Using the PP1-binding mutant PNUTS^W401A^ we were able to determine that the role of PNUTS in endothelial function is mediated via PP1. RNA-sequencing shows how PNUTS loss leads to a dysregulation of gene expression, particularly in axon guidance genes. Finally, we show that aberrant SEMA3B expression is the main mediator of the loss of barrier function after PNUTS depletion.

## METHODS

### Cell culture and cellular assays

Human umbilical vein endothelial cells (HUVEC) were purchased from Lonza and cultured in endothelial cell medium containing supplements and 5% FBS (ECM; ScienceCell). Cells were cultured at 37° C with 5% CO_2_. HUVECs were transfected with control or PNUTS siRNA for 48h. For proliferation assays, cells were incubated with 10 μM EdU for 4 h using the Click-iT EdU microplate Kit (Invitrogen) according to the manufacturer’s protocol. Apoptosis was assessed by incubating the cells with 200nM Staurosporin or medium for 4 h and the caspase 3/7 activity was assayed using the ApoOne Caspase 3/7 Assay (Promega). Senescence associated β-Galactosidase activity was analyzed with the Senescence Associated β-Galactosidase Staining Kit (Cell Signalling Technologies). Images were taken with a bright-field microscope (Axio Observer Z1.0 microscope, Zeiss) and the number of total cells, as well as the number of stained cells was determined in 4 images for per condition and experiment.

### Transfection and lentiviral overexpression

HUVECs were transfected with siRNAs (10nmol/l) with lipofectamine RNAiMax (Life Technologies, Carlsbad, CA), as described before [30]. The siRNAs used were siPNUTS (Sigma-Aldrich Hs01_00133264), siSEMA3B (Dharmacon, J-007754-09-0005) and siMYC (Dharmacon L-003282-02-0005), siRNA Universal Control 1 (Sigma-Aldrich SIC001) was used as control. Lentiviral overexpression of PNUTS was achieved by cloning the full-length human cDNA of PNUTS into pLenti4/v5-DEST (Life technologies). The PP1 non-binding mutant PNUTS (W401A) was kindly provided by Dr. Thomas Küntziger [19] and cloned into pLenti4/v5. All these PNUTS sequences were modified by directed mutagenesis to introduce 3 silent mutations in the seed sequence of the siRNA against PNUTS. pLenti-CMV-MYC was provided by Dr. Linda Penn [27]. Lentiviral particles were generated as described earlier [31].

### Sprouting angiogenesis and immunofluorescence

Endothelial angiogenesis was studied by spheroid sprouting assay. HUVECs were transfected with the indicated siRNAs for 24h. Cells were trypsinized and added to a mixture of culture medium and methylcellulose (80%:20%) and transferred to a 96-well plate to allow the formation of spheroids for 24h at 37°C. The spheroids were collected, added to methylcellulose with FBS (80%:20%) and embedded in a collagen type I (BD Biosciences) gel. Following incubation for 24h at 37°C with 50 ng/ml VEGF the gels were fixed with formaldehyde and microscope images were taken at 10x magnification (Axio Observer Z1.0 microscope, Zeiss). The cumulative length of sprouts was quantified using image analysis software (AxioVision 4.8, Zeiss). For immunofluorescence (IF), 48h after transfection cells were fixed with PFA 4% and incubated with anti-CD31 (1:40, BD Biosciences 550389) or anti-VE-Cadherin (1:400, CST 2500XP), and actin fibers were stained with Acti-stain 555 (Cytoskeleton Inc). Images were obtained in a Nikon A1R confocal microscope. Endothelial gaps were measured by quantifying the intercellular area versus the total area in 4 fields per image, 3 images per experiment.

### Endothelial cell impedance and transwell assays

After lentiviral transduction and/or transfection or stimulation with Tautomycetin (166 nM, R&D Systems), HUVECs were plated in 96-well arrays (96W10idf, Applied Biophysics) at 40,000 cells/well and endothelial cell impedance was measured using the multi-frequency mode for 48 h (ECIS Zθ, Applied Biophysics). Endothelial multifrequency resistance was modeled for the calculation of cell-cell adhesion (R_b_). Transwell assay was performed as explained elsewhere [32].

### Flow cytometry

Cell surface expression of VE-Cadherin and PECAM-1 was analyzed in HUVECs by flow cytometry. Briefly, transfected HUVECs were detached with Accutase and washed in cold incubation buffer (0.1% BSA in PBS). Cells were blocked (5% BSA in PBS) for 30 min on ice and subsequently incubated with fluorophore-labeled antibodies for 30 min on ice. Protein cell surface expression was analyzed using a FACSCalibur™ device (BD Biosciences).

### qRT-PCR and RNA Sequencing

Total RNA from cultured cells was isolated with Direct-zol RNA miniprep (Zymogen) and from tissues using miRNeasy Mini kit (Qiagen) according to the manufacturer’s protocols. Intima RNA was isolated by flushing 200μl of Trizol through the lumen of isolated aortas. Total RNA was reverse transcribed using random hexamer primers and iScript cDNA synthesis kit (BioRad). qPCR was performed with SYBR Green Master Mix on CFX384 Real Time PCR Detection System (BioRad). Gene expression analysis was done using the 2^−ΔCT^ method. RNA obtained from HUVECs and mouse lung tissue was sequenced by Exiqon (whole transcriptome sequencing, 50 bp, 30M reads) and data analysis was performed using DAVID functional annotation tools and Gene Set Enrichment Analysis.

### Western blot

Total cell lysates were collected 48 h after HUVEC transfection or 5 days after lentiviral treatment and western blot (WB) carried as described before [33]. The primary antibodies used were anti-PNUTS (R&D Systems, AF21581), anti-CD31 (Invitrogen, 37-0700), anti-VE-Cadherin (Sigma-Aldrich, V1514), anti-MYC (CST, 9402), anti-phospho-MYC-T58 (abcam, ab185655), anti-tubulin (Thermo Fisher, RB-9281-P) and anti-β-actin (Sigma-Aldrich, A-5441).

### Polysome profiling

siControl and siPNUTS-transfected cells were trypsinized, pelleted and snap-frozen. Polysome profiling was performed as previously described [34]. Briefly, cell pellets were lysed in polysome lysis buffer (gradient buffer containing 100 mM KCl, 10 mM MgCl2, 0.1% NP-40, 2 mM DTT, and 40 U/ml RNasin; Promega, Leiden, Netherlands), and onto 17–50% sucrose gradients and ultracentrifuged for 2 h at 40,000 rpm. The gradients were displaced into a UA6 absorbance reader and absorbance was recorded at an OD of 254 nm.

### ELISA

HUVECs were transfected with siRNA in a 6 well plate as described above. The transfection mix was replaced with 1 ml full medium 4 h post transfection. Supernatants were collected 72 h post transfection and spun down (1000 x *g*, 20 min at 4 °C) to eliminate cell debris. In order to quantify SEMA3B levels, 300 μl supernatant was applied to a human SEMA3B ELISA plate (DLdevelop, Shuigoutou, China), the SEMA3B protein standard was diluted in full medium. The following procedure was performed according to the manufacturer’s instructions.

### Animal procedures

All mice experiments were carried out in accordance with the principles of laboratory animal care as well as according to the German and Dutch national laws. The studies have been approved by the local ethical committees and performed in accordance with the ethical standards laid down in the Declaration of Helsinki. PNUTS^flox/flox^ mice were generated by inserting LoxP sequences, flanking *Ppp1r10* exons 9 to 14, at 69bp downstream of exon 8 and 139 bp upstream exon 15 (Genoway, France). PNUTS^flox/flox^ mice were crossed with the Cdh5-CreERT2 line to obtain the endothelial-specific inducible PNUTS knock-out strain. Cdh5-CreERT2xPnuts^flox/flox^ (PNUTS^EC-KO^) and Pnuts^flox/flox^ (WT) littermates were injected intraperitoneally with 2mg/day tamoxifen dissolved in peanut oil (Sigma-Aldrich) for 5 consecutive days and on the 8^th^ and 10^th^ day after the first injection, and then sacrificed for organ harvest at day 15^th^. Aortic rings were obtained as described [35] and incubated for 48 hours with 10ng/μl VEGF. After fixation, images were taken at 5x magnification and stitched with a Zeiss Axiovert 100 and the endothelial sprouts length were quantified with an image analysis software (AxioVision 4.8, Zeiss). The Evans Blue Extravasation assay was performed on day 14^th^ adapting the protocol by Radu et al 2013 [36]. Briefly, PNUTS^EC-KO^ and WT mice were injected in the tail vein with 25 mg/Kg of Evans Blue solution and observed for 1 h before euthanasia. Then, organs were harvested and incubated for 48 h in formamide to extract the extravasated dye, which was quantified in a spectrophotometer against a standard curve of known concentrations of Evans Blue.

### Histology

Organs were fixed in formalin and embedded in paraffin. 4μm sections were obtained and stained with Periodic-Acid Schiff (Sigma-Aldrich, 395B) and hematoxylin-eosin. Images were obtained in a Leica microscope. Kidney sections were antigen retrieved with citrate buffer and immunostained with an anti-CD31 antibody (1:20, Dianova, DIA-310). Images were taken in a Nikon A1R confocal microscope.

### Statistics

GraphPad 7 (GraphPad Software) was used for statistical analyses. Comparison of two different conditions was analyzed by two-tailed Student’s t test or Mann–Whitney, multiple comparisons were performed by one-way ANOVA using Dunnett’s, Bonferroni or Tukey’s correction. Data are expressed as means ±SEM, p<0.05 was considered as statistically significant (*p<0.05; **p<0.01; ***p< 0.001). The sample size n states the number of independent experiments, unless denoted otherwise. All results were reproduced in at least three technically independent replicates.

### Primers

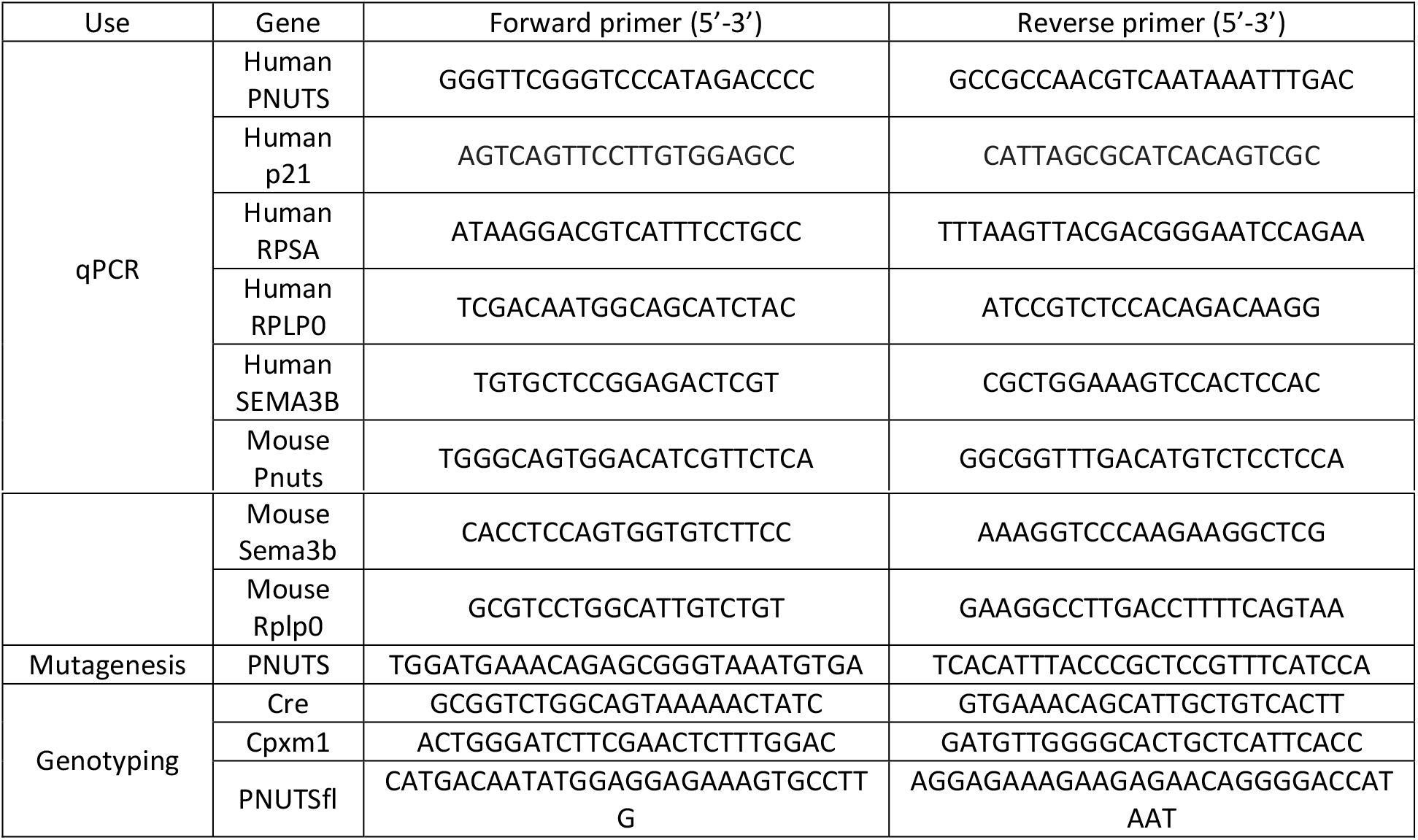

## RESULTS

### PNUTS KD induces senescence of endothelial cells *in vitro*

The first question we sought to answer was whether PNUTS is repressed during aging in the endothelium, as we had described previously for the myocardium, where PNUTS regulates telomere length through TRF2, one of its binding partners [22]. Indeed, PNUTS expression was repressed by 50% in culture-induced senescent HUVECs (Fig 1A). Therefore, we hypothesized that PNUTS may have a role in maintaining endothelial homeostasis which is lost during aging. For this purpose, we used an siRNA-based knock-down (KD) strategy in HUVECs. We selected an siRNA against PNUTS which caused a decrease of PNUTS protein levels of nearly 90% (Fig 1B). Loss of PNUTS provoked an increase in the expression of p21 and in β-galactosidase activity, both markers of senescence (Fig 1C, D). Next to senescence, aging can be accompanied by an increase in apoptosis and a decrease in proliferation. Accordingly, PNUTS KD significantly decreased proliferation (Fig 1E), while apoptosis was not affected. However, PNUTS KD increased cell susceptibility to a pro-apoptotic stimulus (Fig S1), one of the hallmarks of endothelial aging [37]. These results indicate a role for PNUTS in inhibition of endothelial cell senescence and aging.

**Figure 1.**
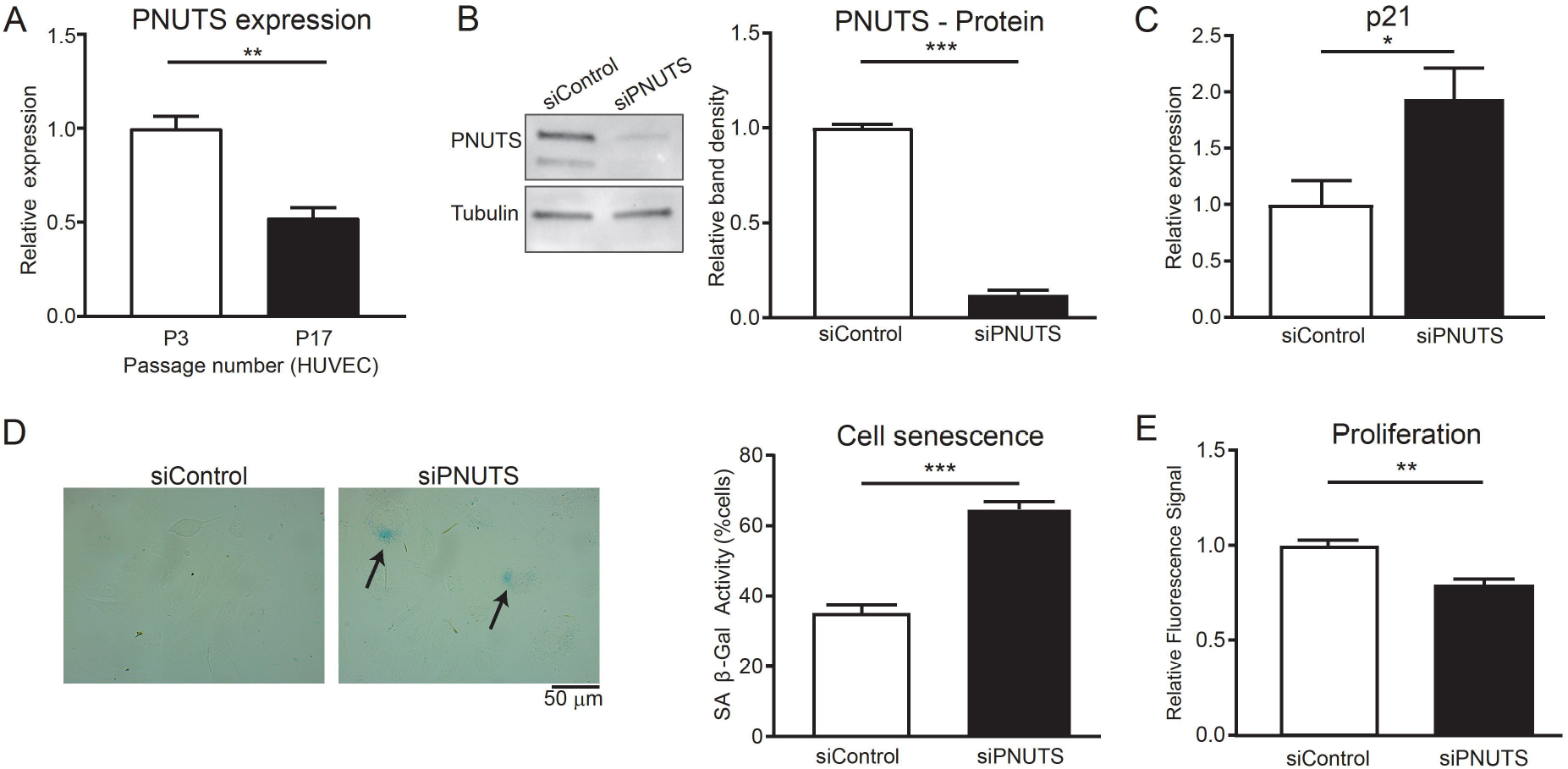
PNUTS is an aging-regulated gene and regulates senescence in endothelial cells. A) HUVECs were cultured for 3 or 17 passages (P) and PNUTS expression was measured by qRT-PCR (n=4) B-E) HUVECs were transfected with siRNAs targeting PNUTS or a control sequence. B) PNUTS protein levels were determined by WB at 48 h after transfection, relative to alpha-tubulin. Densitometric quantification is depicted on the right (n=7). C) Changes in p21 expression was assessed by qRT-PCR. Expression values are relative siControl-treated HUVECs and normalized to RPSA mRNA (n=6). D) The percentage of senescent HUVECs was analyzed by staining for senescence-associated β-Galactosidase (SA-(β-Gal) 72 h after transfection. Images were taken (4 fields per sample) and the percentage of senescent cells (blue) over the total number of cells in each field was calculated in n=3 independent experiments, 2-3 biological replicates per group and experiment). Results are represented as average percentage ± SEM. E) Cell proliferation was assayed by testing the incorporation of EdU after transfection (n=4). *p<0.05, **p<0.01, ***p<0.001.

### PNUTS is necessary for endothelial cell sprouting *in vitro* and *in vivo*

As angiogenesis is typically reduced during aging [37], we performed a spheroid sprouting assay to check the angiogenic capability of PNUTS KD HUVECs. The results indicate that the total length of sprouts, from the surface of the spheroid to the tip cell, was similar in control and KD cells, but in the latter the sprouts showed an aberrant pattern characterized by numerous discontinuities (Fig 2A, B). These results suggest that loss of PNUTS induces loss of stalk cell function or cell-cell adhesions.

**Figure 2.**
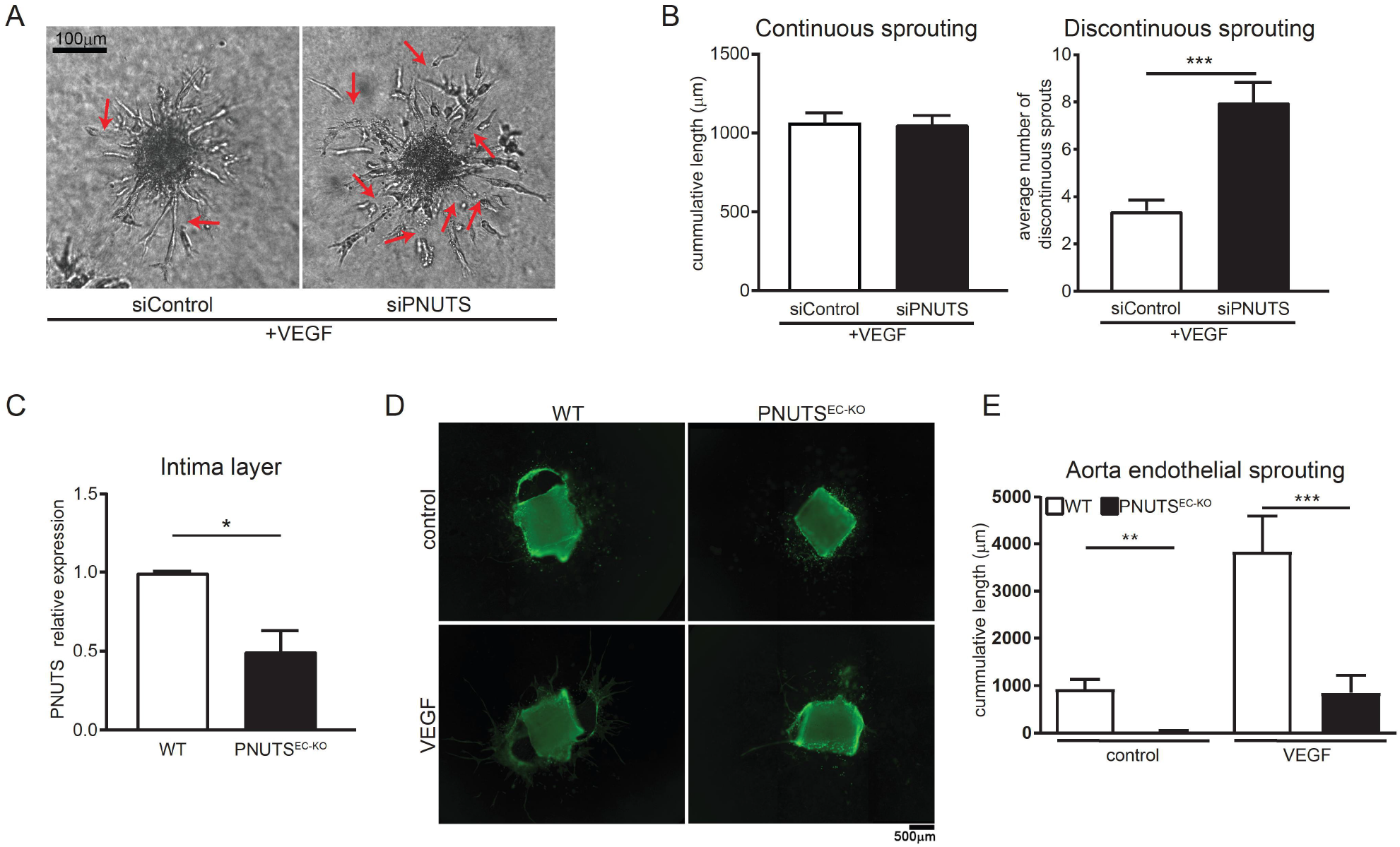
PNUTS is necessary for normal angiogenic sprouting. A) *In vitro* sprouting was analyzed under VEGF (50 ng/ml) stimulation. Representative images are shown, red arrows indicate discontinuous sprouts. B) Quantification of cumulative sprout length (left) and number of discontinuous sprouts (right) (n=30 spheroids per group in 3 independent experiments were measured). C) Endothelial *Pnuts* expression of PNUTS^EC-KO^ and WT mice was assessed by RT-qPCR in intima samples 15 days after initiate tamoxifen treatment and normalized to Rplp0 mRNA (n=3) D) *In vivo* angiogenesis was tested by aortic ring assay in in WT and PNUTS^EC-KO^ mice in basal conditions or upon 48h of VEGF (10 ng/ml) stimulation. E) Quantification of aortic ring sprouting. n=4-5 mice per group *p<0.05, **p<0.01, ***p<0.001.

To assess the physiological role of PNUTS in ECs we sought to use an *in vivo* model of PNUTS loss. To this moment, no loss-of-function mouse model had been described for PNUTS, therefore we generated an endothelial-specific inducible PNUTS knockout mouse line, Cdh5-CreERT2xPnuts^flox/flox^ (PNUTS^EC-KO^), which undergoes efficient loss of PNUTS in the endothelial layer two weeks after tamoxifen treatment compared to PNUTS^f/f^ (WT) (Fig 2C, Fig S2A, B). Using this mouse model, we assessed the angiogenic capability of ECs lacking PNUTS. PNUTS^EC-KO^ aortas were subjected to an aortic ring assay, which showed that the sprouting of aortic rings in PNUTS^EC-KO^ mice was abrogated compared to WT (Fig 2D, E), confirming the findings obtained *in vitro* and suggesting a role for PNUTS in maintaining endothelial cell function.

### PNUTS EC-knockout mice present multiorgan failure and vascular leakage

We carefully monitored PNUTS^EC-KO^ mice after initiation of PNUTS excision by tamoxifen treatment to assess the physiological role of endothelial PNUTS. Within 15 days after tamoxifen treatment, all PNUTS^EC-KO^ mice died, accompanied by a severe accumulation of fluids in peritoneal and pleural cavities. Pathological analysis of kidneys, lungs and hearts showed that PNUTS^EC-KO^ mice suffered from glomerulosclerosis, capillary dilation and pulmonary and cardiac edema (Fig 3A). As our findings pointed that the loss of PNUTS induced a major change in the phenotypical characteristics of ECs, we we assessed global gene expression by RNA sequencing (RNA-seq) in lungs of PNUTS^EC-KO^ and WT mice (Fig 3B, Table S1). The RNA-seq data showed an increase in inflammatory response and coagulation pathways and changes in genes related to endothelial cell-cell and cell-matrix interaction. Intriguingly, the endothelial cell markers *Pecam1* and *VE-Cadherin* were unaltered, pointing to a decreased endothelial cell function rather than a loss of endothelial cells (Fig 3C). Presence of ECs in PNUTS^EC-KO^ kidneys was confirmed by IF of PECAM1 in renal glomeruli, although the distribution of PECAM1 in PNUTS^EC-KO^ glomeruli suggests a loss of capillary organization (Fig 3D). Intriguingly, the phenotype observed in several organs, including the presence of fluid in the cavities, pointed to increased microvascular permeability. Therefore, we investigated whether PNUTS^EC-KO^ mice suffered from vascular leakage. We performed Evans Blue extravasation assays and found that the tracer was able to cross the endothelial barrier to the tissue in several organs in PNUTS^EC-KO^ mice, as well as the peritoneal cavity, pointing to a loss of barrier integrity *in vivo* (Fig 3E,F). These results suggest that endothelial PNUTS is critical for maintenance of endothelial barrier and its loss is incompatible with life due to severe vascular leakage.

**Figure 3.**
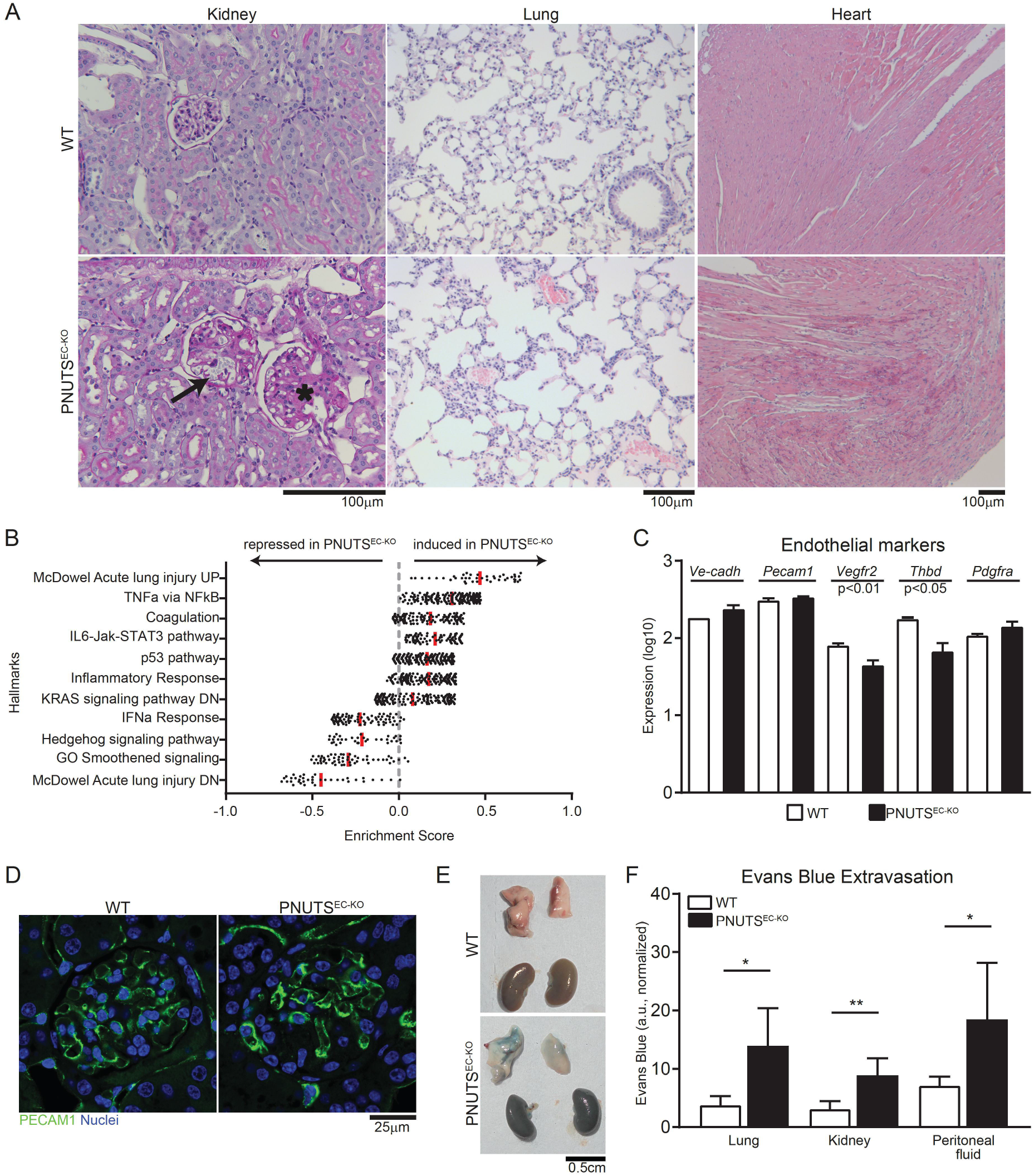
Induction of endothelial-specific PNUTS KO in mice provokes multiorgan failure due to vascular leakage. A) Histopathological study of different tissues in PNUTS^EC-KO^ mice compared to WT mice: left, kidney samples (Periodic-Acid Schiff staining) showing glomerulosclerosis (asterisk) and capillary dilatation (black arrow) in renal glomeruli; middle, lung samples (haematoxylin-eosin) presenting thrombi; right, heart samples (haematoxylin-eosin) presenting edema. B) RNA-seq was performed with lung tissue of WT and PNUTS^EC-KO^ mice and was analysed for differentially regulated pathways using Gene Set Enrichment Analysis. Enrichment scores of the indicated pathways are plotted on the *x* axis (n=3). C) The expression levels of the entohelial markers *Ve-cadherin, Pecam1, Vegfr2, Thbd* and *Pdgra* in WT and PNUTS^EC-KO^ lung samples was confirmed by RT-qPCR and normalized to *Rplp0* mRNA (n=3). D) The presence of ECs in the glomerular capillary network was investigated by PECAM1 immunostaning in kidneys of WT and PNUTS^EC-KO^ mice. E-F) An Evans Blue (EB) extravasation assay was performed to measure the vascular extravasation in different organs. E) Representative lung and kidney images 1 h after intravenous administration of EB at 25 mg/kg. F) Colorimetric measurement of extravasated EB into lung, kidney and peritoneal fluid (n=4-5 mice per group, *p<0.05, **p<0.01).

### PNUTS loss compromises the endothelial barrier by disrupting cell-cell interaction in a PP1-dependent manner

The endothelial monolayer is responsible for regulating the trafficking of solutes and cells from the blood to the surrounding tissues, forming a selective barrier between both. In light of our *in vivo* results, we confirmed whether PNUTS is necessary for this function. We measured the impedance of HUVEC monolayers using an electrical cell impedance system (ECIS) and found that loss of PNUTS decreased the resistance of the endothelial monolayer (Fig 4A, B), suggesting an increase in endothelial permeability. As a confirmation of these results, we performed a transwell assay, which showed an increased leakage through the monolayer of PNUTS-silenced cells, as measured by passage of HRP (Fig 4C). This decrease in endothelial barrier function is mainly due to a loss of endothelial cell-cell interaction, as indicated by multifrequency modelling of R_b_ (Fig 4D, E). IF analysis as well as Western blot and flow cytometry measurements of HUVECs showed that the effects of PNUTS depletion on endothelial barrier function are not due to decreased levels or delocalization of adherens junction proteins (Fig 4F, G and S3). However, there is an increase in the presence of intercellular gaps that might explain the inability to form an efficient barrier in the absence of PNUTS (Fig 4H, S3). Altogether, this suggests that PNUTS is necessary to maintain a functional endothelial monolayer through a mechanism independent of adherens junction protein expression.

**Figure 4.**
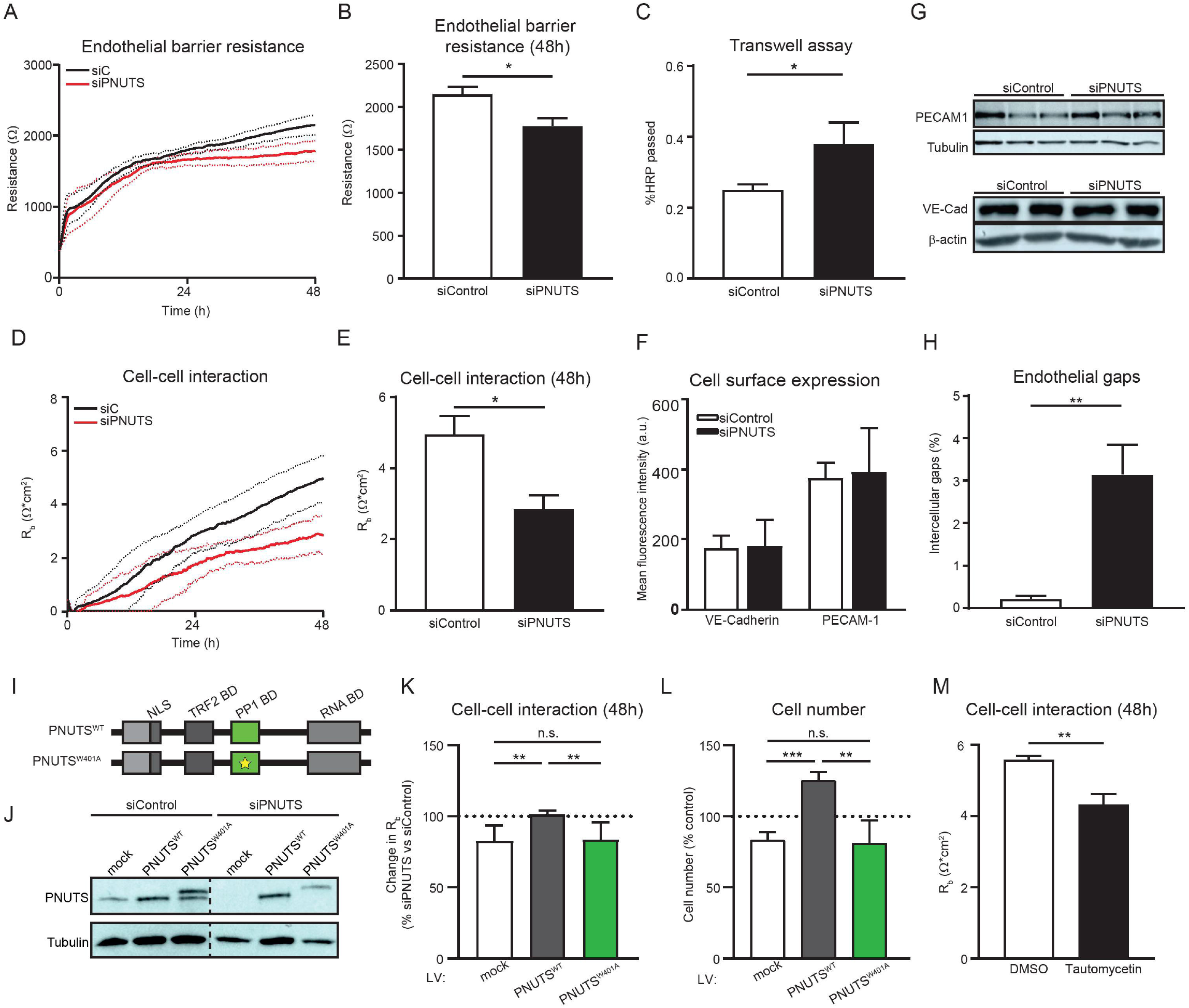
PNUTS is necessary for endothelial barrier function independently of cell junction gene expression. A-H) HUVECs were treated with siRNA (si) targeting PNUTS or a control sequence. A-B) HUVEC barrier resistance was assessed by ECIS at 4000 Hz for 48 h, presented as average resistance ± SEM of n=3 experiments, 8 biological replicates per group and experiment. C) HUVECs were seeded in transwells and HRP passage through the endothelial monolayer was assessed by absorption measurements (450 nm) and shown as percentage of total HRP (n = 3). D-E) Cell-cell interaction was assessed by modelling of data obtained by ECIS measured as R_b_ (Ω*cm^2^); presented as average resistance ± SEM of n=3 experiments, 8 biological replicates per group and experiment. F) Cell surface presence of PECAM-1 and VE-cadherin was assessed by flow cytometry 48 h after siRNA transfection (n=3). G) Total PECAM-1 and VE-cadherin levels were assessed by Western Blotting (WB). Tubulin and β-actin expression was used as loading control. H) Cells were grown into confluence and immunostained for VE-cadherin, PECAM-1 and F-actin. DAPI was used to stain nuclei. The presence of intercellular gaps in endothelial monolayers was measured by quantifying the intercellular areas versus the total area in 4 fields per image, 3 images per experiment, n=4. I) Schematic representation of the PNUTS lentiviral vectors used for the barrier rescue experiments. Both vectors included silent mutations in the seed sequence of siPNUTS. HUVECs were transduced with the indicated constructs for 8-10 days and later transfected with siControl or siPNUTS to silence endogenous expression of PNUTS, before subjecting them to ECIS and cell counting (4-6 independent experiments, 4 biological replicates per group and experiment). J) PNUTS expression in total cell lysates of HUVECs treated with indicated vectors and siRNAs was analyzed by WB, Tubulin was used as a loading control. Representative WB is shown. K) Change in cell-cell interaction measurement from ECIS 48 h after cell seeding, measured as variation of R_b_ of siPNUTS-vs siControl-treated cells. L) Cell proliferation of ECIS-assayed HUVECs, measured as % number of cells relative to control. M) HUVECs were treated with vehicle or Tautomycetin (166 nM) and cell-cell interaction was measured for 48 h by ECIS 9n=3). *p<0.05, **p<0.01, ***p<0.001.

Previous studies describe PNUTS as a binding partner and activator for PP1. We therefore aimed to assess the potential role of PP1 for PNUTS function by using several lentiviral vectors to overexpress siRNA-resistant wild-type (PNUTS^WT^) and PP1-non-binding mutant PNUTS (PNUTS^W401A^) (Fig 4I). All of them proved to be resistant to siPNUTS treatment (Fig 4J) and were used to test the relevance of the PP1 binding domain for the role of PNUTS in barrier maintenance by ECIS. As seen in Fig 4K, only the overexpression of PNUTS^WT^ was able to rescue the drop in barrier function caused by PNUTS KD, while the PP1-binding mutant was not. Moreover, the PP1-binding is important for endothelial survival, as the overexpression of the PNUTS^W401A^ mutant decreased EC proliferation to the same levels as siPNUTS treatment, while PNUTS^WT^ had a positive effect on proliferation of ECs (Fig 4L). Furthermore, the role of PP1 activity in endothelial barrier maintenance was assessed. We used Tautomycetin, a PP1-inhibiting drug, in HUVECs and analysed cell-cell interactions by ECIS. PP1 inhibition triggers a loss of barrier function in the endothelial monolayer (Fig 4M), suggesting that the PNUTS-PP1 complex is a critical mediator for endothelial barrier maintenance.

### PNUTS loss triggers a complex transcriptional profile change in endothelial cells

In order to more elaborately study the mechanism by which PNUTS controls EC function, we studied the gene profile elicited by PNUTS loss. We performed RNA-seq of si-Control and siPNUTS-treated HUVECs. We found 7034 transcripts significantly up- or downregulated (Table S2). KEGG pathway functional annotation identified several affected pathways, among which mRNAs encoding ribosomal proteins stood out (Fig S4A). Sixty-eight ribosomal protein-encoding genes were significantly altered, of which 62 were strongly downregulated (Fig S4B). In line with this finding, a polysome profiling of siControl- and siPNUTS-treated HUVECs showed that PNUTS KD provokes a reduction of ribosomal content in ECs (Fig S4C).

Gene set enrichment analysis showed a decrease in expression of Myc-dependent targets (Fig S5). MYC has been described to have roles in endothelial proliferation and angiogenesis [38] and it has been recently suggested that Myc phosphorylation and subsequent degradation might be controlled by the PNUTS-PP1 axis [27]. MYC protein levels were decreased in PNUTS KD cells, while phosphorylation of MYC at Threonine 58 was increased, which induces ubiquitination and proteasomal degradation of MYC (Fig S5C). However, siRNA-mediated KD of MYC had no effect on endothelial barrier function, while lentiviral-mediated overexpression of MYC had a negative effect, opposite to expected (Fig S5D). Therefore, MYC is a negative regulator of the endothelial barrier in a PNUTS-independent manner.

### PNUTS regulates barrier function via suppression of SEMA3B expression

The complexity of the data obtained by RNA-seq led us to compare the gene expression profile after PNUTS loss in HUVECs and mouse lungs. A total of 277 common genes were found to be regulated in both datasets. KEGG pathway analysis of these revealed a significant presence of axon guidance pathway genes (Fig 5A). We confirmed the RNA-seq data by qPCR in PNUTS-silenced HUVECs, finding a dramatic induction of SEMA3B expression (Fig 5B), one of the highest upregulated genes after PNUTS KD (Fig 5C, Table S2). SEMA3B protein was also found to be increased in the supernatant of siPNUTS-treated cells (Fig 5D), as quantified by ELISA. Furthermore, the expression of *Sema3b* in the aortic intima of PNUTS^EC-KO^ mice is increased 7-fold compared to WT mice (Fig 5E). SEMA3B has been described previously as a secreted protein that causes repulsion signals in ECs which interferes with correct angiogenesis [39], but has not yet been linked to endothelial barrier formation.

**Figure 5.**
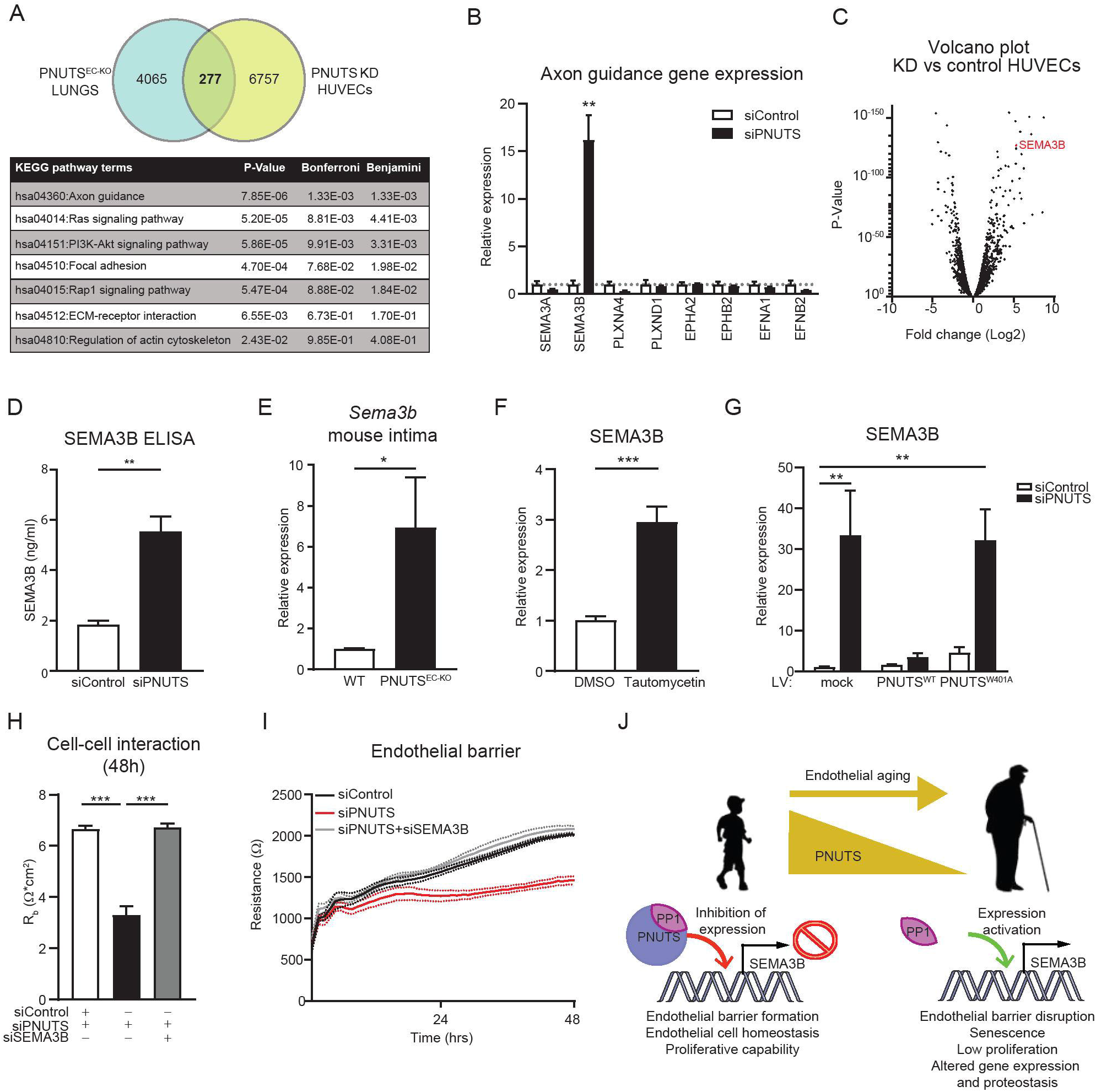
PNUTS KD induces transcriptomic changes in HUVECs. A) The two subsets of RNAseq data we obtained, from endothelial-depleted PNUTS mouse lungs and PNUTS knockdown (KD) HUVECs, were compared to find common targets of PNUTS depletion. The Venn diagram depicts the finding of 277 common transcripts, which were functionally analysed using KEGG pathways analysis, shown in the table below. B) Changes in axon guidance gene expression after PNUTS silencing was assessed by RT-qPCR. Expression values are relative siControl-treated HUVECs and normalized to RPSA mRNA (n=6). C) Volcano plot showing the distribution of gene expression in PNUTS KD versus control HUVECs. SEMA3B is marked in red. D) Supernatant SEMA3B concentration was determined by ELISA 72 h after PNUTS silencing (n=4). E) Expression levels of *Sema3b* mRNA in intima samples of PNUTS^EC-KO^ mice was assessed by RT-qPCR, relative to WT samples and normalized to *Rplp0* mRNA. F) HUVECs were treated with vehicle or Tautomycetin (166 nM) for 48 h and mRNA was analyzed by RT-qPCR for expression of SEMA3B, normalized to *RPSA* mRNA (n=3). G) Expression of SEMA3B was measured in mRNA samples of HUVECs assayed in Fig 4 J-L, relative to control cells and normalized to *RPSA* mRNA H-I) HUVECs were co-transfected with siPNUTS and/or siSEMA3B were subjected to ECIS for 48h (n=4 independent experiments, 4 biological replicates per group and experiment) *p<0.05, **p<0.01, ***p<0.001 H) Cellcell interaction was modeled. I) Endothelial resistance was measured at 4000 Hz. J) Graphic summary of the proposed mechanism. In young individuals, PNUTS interacts and promote activity of PP1, which represses the expression of SEMA3B. Endothelial cells are in homeostasis and maintain their barrier function. During aging, PNUTS is repressed in endothelial cells. The absence of PNUTS inhibit PP1 function at the SEMA3B promoter activating SEMA3B expression. SEMA3B exerts repulsive signals between endothelial cells, promoting intercellular gaps and disrupting the barrier. This provokes a series of critical changes in the cells leading to cellular senescence.

PP1 has been described to exert a broad role in regulating gene expression, thus we studied SEMA3B expression after treating HUVECs with the PP1 inhibitor drug Tautomycetin. Indeed, PP1 inhibition triggered a significant induction of SEMA3B expression (Fig 5F). Furthermore, the induction of SEMA3B expression after PNUTS silencing is rescued by overexpression of PNUTS^WT^, but not by overexpression of the PP1-binding mutant PNUTS^W401A^ (Fig 5G). Finally, we hypothesized that SEMA3B might be the causal mechanism triggered by loss of PNUTS leading to a disruption of barrier function. To test this, we co-treated HUVECs with siPNUTS and siSEMA3B and found that SEMA3B knock-down completely abolished the effect of PNUTS loss on cell-cell interaction and endothelial resistance (Fig 5H-I). Altogether, these results points to the axon repulsion signal SEMA3B as a disruptor of endothelial monolayer integrity which is repressed by PNUTS in homeostatic situations through PP1 binding and activity.

## DISCUSSION

This study describes the aging-regulated protein PNUTS as a critical element to maintain endothelial homeostasis, as loss of endothelial PNUTS elicits a phenotype of endothelial leakage compromising homeostasis. PNUTS regulates PP1 function in the endothelium, acting as a binding platform for the enzyme and conforming an axis that is necessary for barrier integrity. Loss of PNUTS or PP1 activity deregulates the signature gene expression profile in ECs and triggers the induction of SEMA3B, a member of the axon guidance family of proteins that acts as a repulsion signal for ECs.

PNUTS structure and function have been mainly studied *in vitro* in cell lines, while the role of PNUTS in physiology and disease has been largely unknown. In particular, the role of PNUTS in the cardiovascular system or aging has been described previously only in the myocardium, where PNUTS regulates telomere length through TRF2 in cardiomyocytes and is downregulated during aging or after myocardial infarction [22]. In the present work, we sought to assess the role of PNUTS in the endothelium. As seen in cardiomyocytes, endothelial PNUTS is aging-regulated and critical to maintain cell functionality in ECs. In fact, PNUTS loss recapitulates many of the hallmarks of endothelial aging, such as expression of senescence markers, decreased proliferation, gene expression deregulation and endothelial dysfunction (Fig 1, 2, 4 and S1). However, PNUTS loss in the myocardium provoked an increase in DNA damage and apoptosis through the aforementioned mechanism, whilst in the endothelium PNUTS has a role in maintaining endothelial integrity via PP1-mediated induction of SEMA3B. This suggests that PNUTS function and mechanism of action is cell type- and binding partner-specific. PNUTS was initially described as a binding platform for PP1, and although several other binding partners of PNUTS have been described over time [40], the main body of knowledge on PNUTS function continues to be related to this phosphatase. The PNUTS-PP1 axis has been suggested to be responsible of DNA damage response [19], MYC activation [27], cell cycle entry [21], apoptosis [28], gene expression regulation [26] and transcription [29]. None of these studies report, however, a role for PNUTS-PP1 in physiological processes, with the exception of the work of Ciurciu and colleagues, which modelled their findings in the embryonic development of *D. melanogaster*. Our work is the first, to our knowledge, that studied the contribution of this axis in a physiological context in mammals.

The main function of ECs is to form a barrier between the blood and the underlying tissues that allows correct exchange of nutrients and waste products. This critical function is also impaired by aging, which induces hyperpermeability in the peripheral tissues and eventually edema formation due to a loss in the integrity of adherens junctions [41]. Notably, our results in PNUTS^EC-KO^ mice and PNUTS KD HUVECs point to endothelial barrier dysfunction independent of adherens junctions (Fig 3 A,E,F and 4 A-G), which we concluded was the main consequence of PNUTS loss in the endothelium and that it is mediated by PP1 binding and activation (Fig 4H-M). Previously, PP1 had been described to have a role in endothelial permeability by dephosphorylation of cytoskeleton proteins [42], but here we show that PP1 function in ECs is linked to a nuclear co-factor (PNUTS).

Hence, we considered a nuclear mechanism that provoked cells to be less prone to form proper interactions. PNUTS loss elicits a complex response in gene expression both *in vivo* (Fig 3B) and *in vitro* (Fig S4A, S5A), which led to the identification of SEMA3B as potential target of the PNUTS-PP1 axis. This secreted semaphorin, as others of the axon guidance family of genes, was historically studied for its repulsion role in the guidance of neuronal growth in the developing neural system [13], but new roles in endothelial barrier maintenance, preeclampsia and inhibition of angiogenesis have emerged [15] [43] [39], which aligns with our findings. Little is known about the regulation of SEMA3B expression. Our study suggests that SEMA3B is induced in the absence of a working PNUTS-PP1 complex (Fig 5 B-G), and that it is the main actor in impeding the formation of a healthy endothelial barrier (Fig 5 H,I) upon loss of PNUTS. We hypothesize that PNUTS may restrict SEMA3B expression by PP1-mediated dephosphorylation of epigenetic modulators or transcription factors [26] [29] [44] [45]. Finally, our results highlight a novel concept that explains loss of barrier function during aging in which dropping PNUTS levels result in aberrant activation of SEMA3B expression that induces repulsion of ECs.

In conclusion, this study provides evidence that PNUTS acts as a PP1-binding partner which is critical for endothelial function during aging. Our data highlight the importance of the PNUTS-PP1 axis in providing a tight control of the endothelial gene expression profile, which is necessary to maintain barrier functionality through the regulation of the semaphorin family and signal towards SEMA3B as a potential clinical target against aging-related vascular hyperpermeability.

---

**Supplemental Figure 1.** HUVECs were transfected with siControl and siPNUTS for 48 h and subsequently treated with vehicle or Staurosporin 200 nM. Apoptosis was assayed by determining their Caspase 3/7 activity (n = 5).

**Supplemental Figure 2.** A) Schematic representation of the PNUTS-floxed allele and further recombination of LoxP regions after crossing the PNUTS^fl/fl^ mice with the inducible endothelial-specific Cdh5-CreERT2 strain and treatment with tamoxifen. B) Time course of the experimental work in mice.

**Supplemental Figure 3.** HUVECs were transfected with Control and anti-PNUTS siRNA (si). 72h after transfection, full-grown monolayers were analyzed by IF staining using anti-VE-cadherin (white), anti-PECAM1 (green) and F-actin (red). DAPI (blue) was used to stain nuclei. Representative images are shown.

**Supplemental Figure 4.** HUVECs were transfected with Control and anti-PNUTS siRNA (si) for 48 h. Total RNA was isolated and the transcriptome was studied by RNA-seq. (n=3) A) Functional analysis of RNA-seq data clustered as KEGG pathway terms shows the differentially expressed pathways. B) Heat map of differentially regulated genes encoding for ribosomal proteins upon PNUTS KD. C) Polysome profiling of siControl (black line) and siPNUTS (red line) HUVECs. The consecutive peaks show the presence of 40S and 60S ribosome subunits, 80S ribosome (isolated ribosomes, normally inactive) and polysomes (two or more ribosomes bound to a single mRNA molecule, actively translating into protein) (n=3).

**Supplemental Figure 5.** A) Left, RNA-seq from PNUTS-silenced HUVECs was also analysed for differentially regulated pathways using GSEA. Enrichment scores of the indicated pathways are plotted on the x axis. Right, GSEA of MYC target datasets in PNUTS KD cells. ES, enrichment score; NES, normalized enrichment score. B) Heat map of differentially regulated MYC signature genes upon PNUTS KD. C) Left, representative WB of total cell lysates blotted with anti-MYC and anti-pMYC-T58 antibodies, anti-actin was used as loading control; right, densitometry quantification of MYC bands and pMYC-T58/MYC ratio in siControl and siPNUTS HUVECs. D) HUVECs were transfected with siRNAs against PNUTS and MYC or transduced with a lentiviral vector to overexpress MYC, and endothelial barrier resistance was assessed by ECIS (n=3, 4 biological replicates per group and experiment). *p<0.05, ***p<0.001

