## Supplementary figures and images for "The PNUTS-PP1 axis regulates endothelial aging and barrier function via SEMA3B suppression"

### Supplemental Figure 1

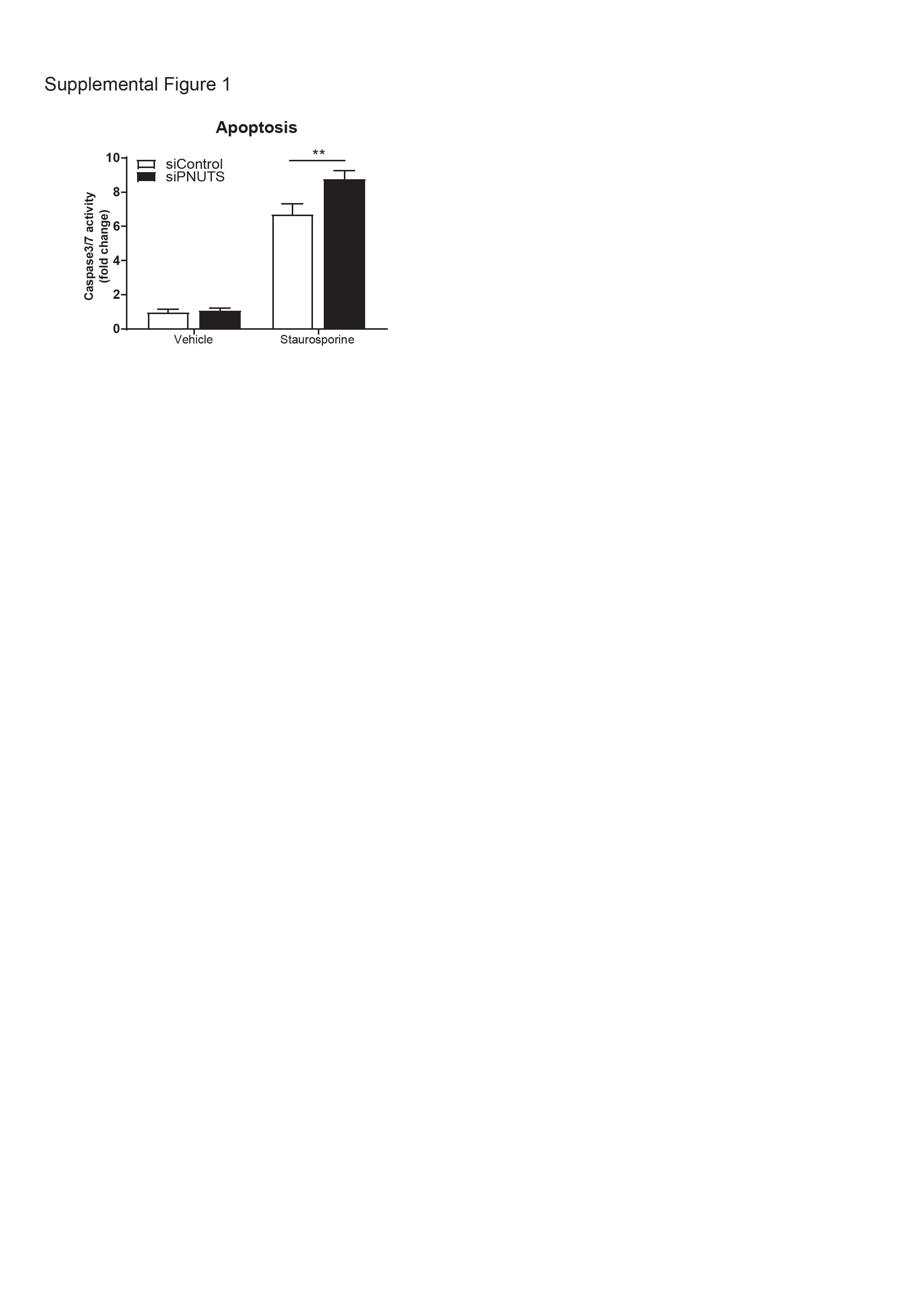

### Supplemental Figure 2

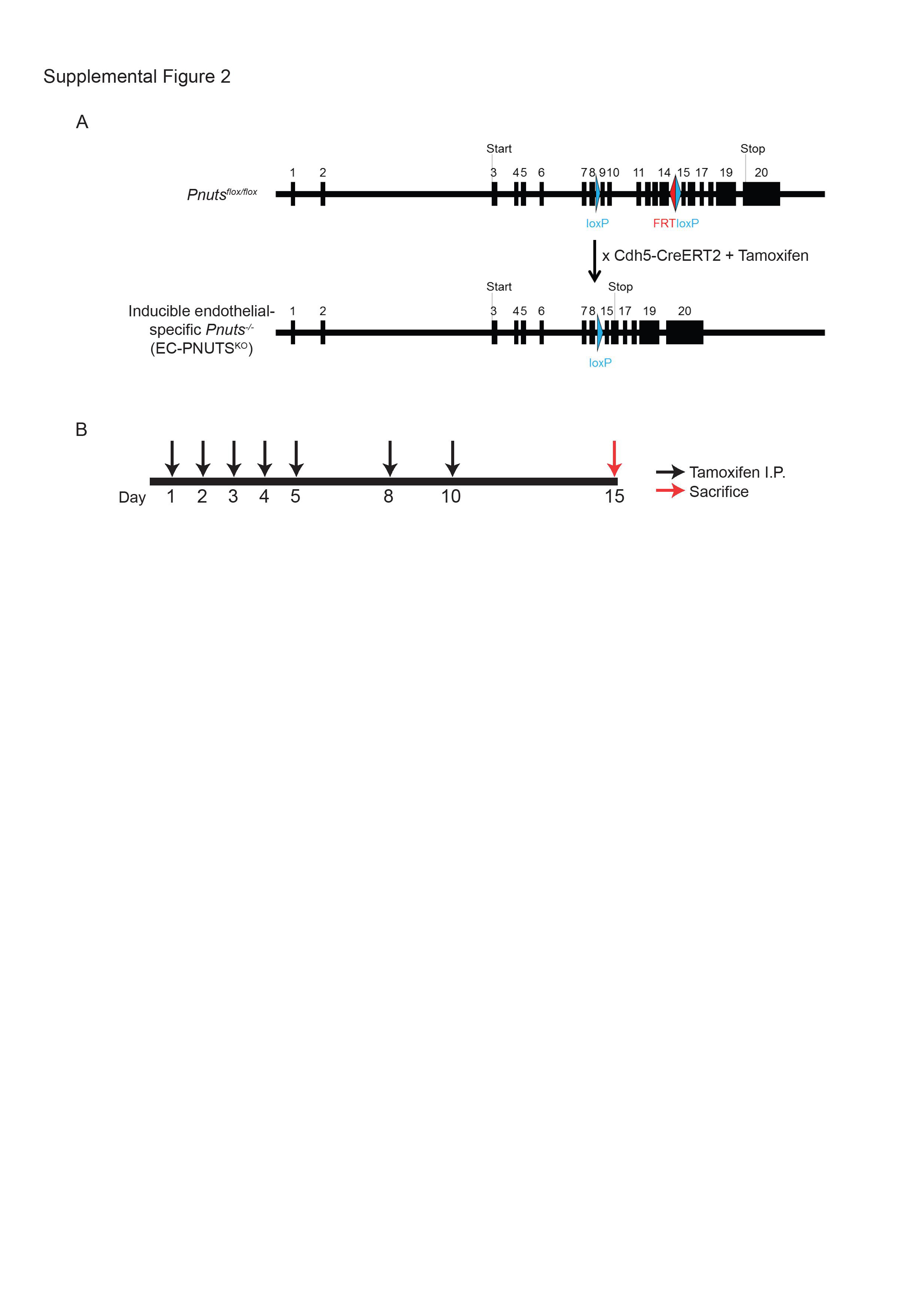

### Supplemental Figure 3

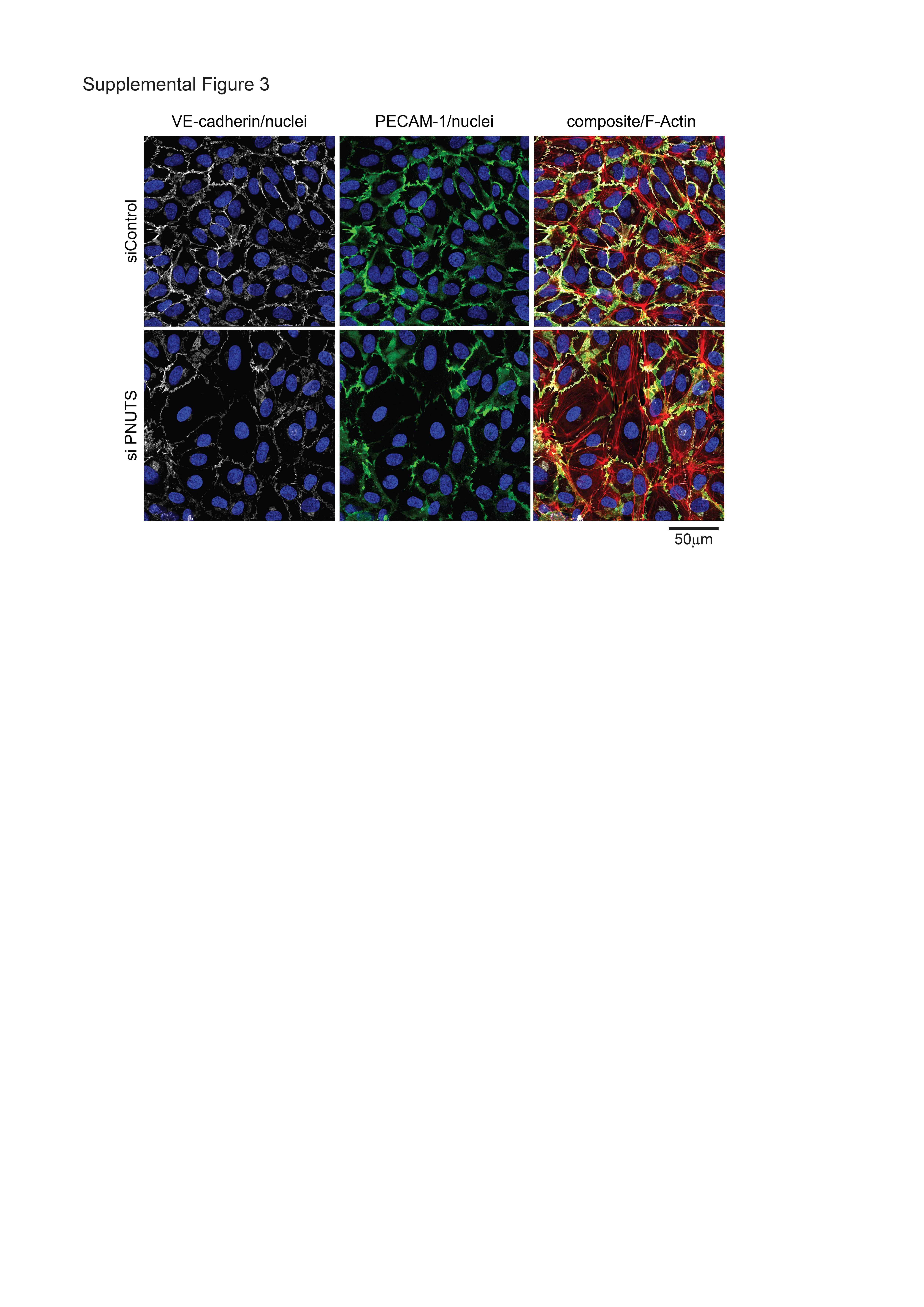

### Supplemental Figure 4

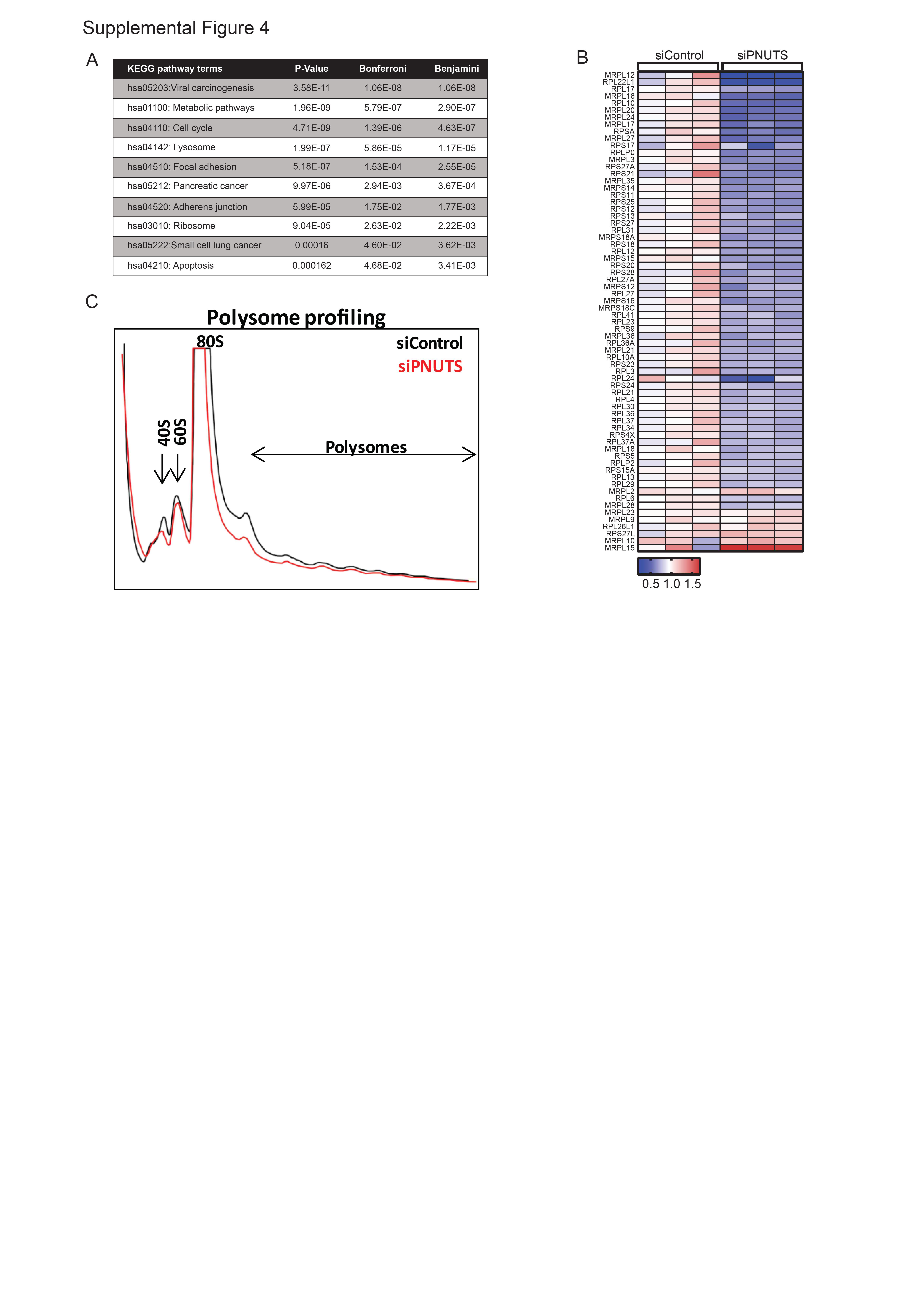

### Supplemental Figure 5

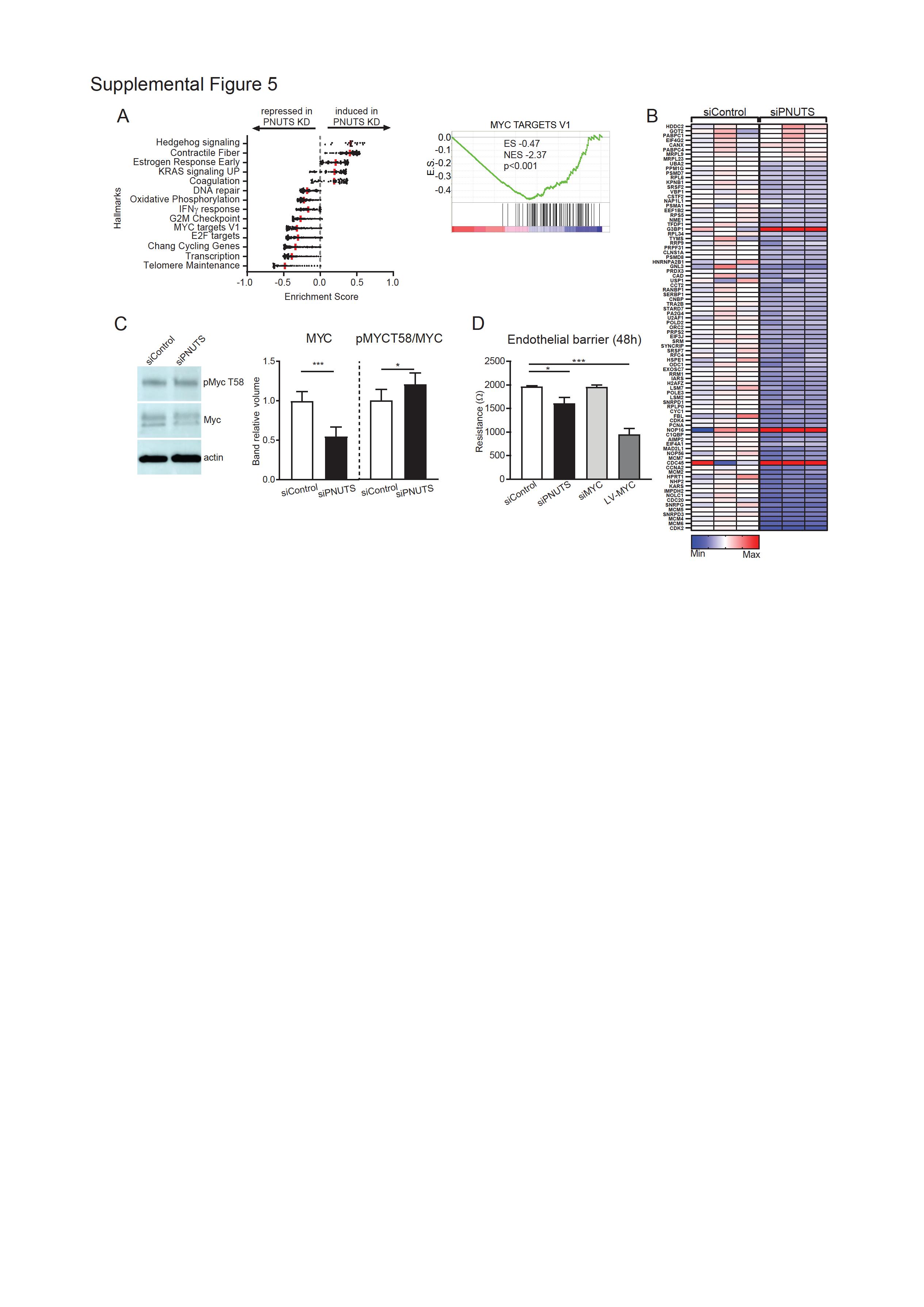
